# CD4+ T-Cells Drive Triple Negative Breast Cancer Recurrence via Non-Canonical TGFβ Signaling

**DOI:** 10.64898/2026.08.14.744926

**Authors:** McKenzie A. Mayeaux, Benjamin P. Altman, Benjamin C. Hacker, Steven M. Alves, Dadi Jiang, Albert C. Koong, Edward E. Graves, Marjan Rafat

## Abstract

Radiation therapy is a cornerstone of breast cancer treatment and reduces recurrence overall. However, patients with triple negative breast cancer (TNBC) continue to experience recurrence at higher rates than patients with other subtypes, especially when immunocompromised. While CD8⁺ T-cells are known to mitigate recurrence, the role of CD4+ T-cell subsets in shaping the irradiated microenvironment remains unclear. We show that irradiated mammary tissue from mice accumulates CD4+ T-cells and exhibits a TGFβ-enriched cytokine milieu coincident with macrophage infiltration. We demonstrate that Th2-polarized CD4+ T-cells promote invasion of TNBC cells and macrophages through secretion of TGFβ. Neutralization of TGFβ significantly reduces this invasive phenotype. Mechanistically, Th2-conditioned media induces *Tgfb1* expression in both TNBC cells and macrophages, establishing a TGFβ-dependent feed-forward amplification loop. In TNBC cells, Th2-derived TGFβ activates non-canonical signaling characterized by increased p38 MAPK and NF-κB phosphorylation, linking cytokine exposure to pro-invasive behavior. Together, these findings identify Th2-derived TGFβ as a driver of pro-invasive tumor reprogramming and suggest that interruption of Th2–TGFβ signaling may prevent recurrence following therapy.

## Introduction

Breast cancer is one of the leading causes of death for women worldwide^1,2^. Triple negative breast cancer (TNBC) only accounts for 15-20% of all diagnosed breast cancers but has disproportionately high mortality and a high rate of fatal recurrence^3,4^. TNBC is characterized by lack of hormone receptors for estrogen (ER) and progesterone (PR) and overexpression of HER2, rendering hormone-targeted therapies ineffective. Because of this, the standard of care for patients with TNBC include surgery, immunotherapy, chemotherapy, and radiation therapy. Most breast cancer patients receive radiation due to its favorable tumor control^5^. However, damage to normal tissue is unavoidable.

While radiation therapy lowers rates of recurrence overall^6^, retrospective studies have shown that sustained lymphopenia, or reduced absolute lymphocyte count ^7^, post-therapy leads to worse outcomes and increased recurrence^8,9^. In mouse models of lymphopenia, circulating tumor cells from breast tumors homed to irradiated mammary tissue. Excess macrophage infiltration preceded tumor cell invasion in immunocompromised mice. Further studies revealed that CD8+ T-cells were protective against such infiltration^9^. However, CD4+ T-cells accumulated but were not protective against macrophage and tumor infiltration.

CD4+ T-cells are known to direct macrophage phenotypes^10^. While Th1 cells can sustain the inflammatory M1 phenotype, Th2 cells support the immunosuppressive, wound-healing M2 tumor-associated macrophages (TAMs), which are further shaped by the CD4+ T-cell secretome and facilitate metastasis^11^. In the context of radiation, CD4+ T-cells favor Th2 programming over Th1^12^, and depleting CD4+ T-cells, blocking Th2 polarization, or neutralizing signaling mediators improve radiation-associated tumor control^11,13^. The clinical significance of immune cell presence in the tumor microenvironment is demonstrated in clinical studies that associate high CD4:CD8 or Th2:Th1 ratios with higher tumor grade, advanced stage, metastasis, and reduced overall survival^14^.

Despite extensive study of tumor-intrinsic immunology, damage to the normal adjacent tissue in influencing recurrence remains unknown. Here, we establish that an irradiated environment enriched for transforming growth factor-beta (TGFβ) increases TNBC cell and macrophage invasive behavior. Furthermore, TGFβ secreted by Th2 cells is vital for p38 expression and NF-κB phosphorylation in TNBC cell while supporting the recruitment of macrophages. We show that a TGFβ1-driven feed-forward loop links Th2 cells, macrophages, inflammation, and immunosuppression to TNBC recurrence. Elucidating the mechanisms that drive or permit TNBC recurrence post-therapy may lead to improved treatment regimes and patient outcomes.

## Methods

### Cell Culture

Luciferase-labeled 4T1 mouse mammary carcinoma cells were obtained from Dr. Christopher Contag (Stanford University) in August 2011 and cultured in in RPMI-1640 (Gibco, Grand Island, NY, USA 11875119). All cell lines tested negative for mycoplasma contamination with the MycoAlert Mycoplasma Detection Kit (Lonza, Basel, Switzerland LT07-318). Cells were used within three passages before injection into mice. All cells were maintained at 37°C in a humified incubator containing 5% CO_2_.

For primary macrophage culture, bone marrow-derived macrophages (BMDMs) were isolated from the femurs of female 8-10 week old BALB/c mice as described previously^15–18^. Isolated cells were cultured with 10 mg/mL macrophage stimulating serum factor (MCSF; Invitrogen, Carlsbad, CA, USA PMC2044) for 7 days on low adhesion plates until maturation into macrophages. BMDMs were cultured in IMDM (Gibco 12440061) supplemented with 10% FBS, antibiotics (PenStrep, Gibco 15140-122), 1% HEPES (Corning MT25060CI), and fresh MCSF (10 ng/mL, Gibco PMC2044). Primary CD4+ T-cells were isolated from the spleens of 8-10 week old female BALB/c mice with the Miltenyi CD4 T-cell Isolation kit (130-104-454). For Th2 polarization, CD4+ T-cells were stimulated with aCD28 (BD, San Jose, CA, USA BDB553295), aCD3 (BD BDB553057), aIFNγ (Invitrogen MM700), and IL-4 (Peprotech, Cranbury, NJ, USA 214-14) for 48 hours and then cultured in Thermofisher Optimizer media (Gibco A1048501) supplemented with L-glutmine, 2-mercaptoethanol (Gibco 21985023), rIL2 (NIH), and PenStrep (Gibco 15140122). Conditioned media (CM) from Th2 cells was collected every 2 days, filtered, and stored at -80°C until use.

### Orthotopic Tumor Injection

Animal studies were performed in accordance with institutional guidelines and protocols approved by the Vanderbilt University Institutional Animal Care and Use Committee. In compliance with ARRIVE 2.0 guidelines^19^, all animal studies were performed on age, sex, and strain matched animals that were treated at the same time of day. Tumor inoculation was performed by injecting 5×10^4^ 4T1 cells in a volume of 50 µL directly into the number 4 right mammary fat pads (MFP) of 8- to 10-week- old female BALB/c mice. In T-cell depletion experiments, 0.5 mg anti-CD4 (GK1.5; BioXCell, Lebanon, NH, USA BE0003-1) and/or 0.5mg anti-CD8a (2.43; BioXCell, BP0061) was injected intraperitoneally every 5 days starting from the day of inoculation. Control mice were injected with 0.5 mg rat IgG2b isotype control (LTF-2; BioXCell BE0090) using the same dosing schedule. All mice were purchased from Charles River Laboratories (Wilmington, MA, USA).

### Radiation

Mouse MFPs were irradiated to 20 Gy using a 250 kVp cabinet x-ray system filtered with 0.5 mm Cu^9^. Mice were anesthetized by administering 80 mg/kg ketamine hydrochloride and 5 mg/kg xylazine intraperitoneally and then shielded using a 3.2 mm lead jig with 1 cm circular apertures to expose MFPs. Transmission through the shield was less than 1%.

### Luminex Multiplex Cytokine Assay

The MFPs of control and irradiated BALB/c mice with or without CD8+ T cell depletion were resected 10 days post-radiation. To assess cytokine profiles at the local site of infiltration, MFPs were harvested and homogenized in 20 mM Tris HCl (pH 7.5) buffer with 0.5% Tween 20, 150 mM NaCl, and protease inhibitor, centrifuged for 10 min at 4 °C, and supernatant was stored at − 80 °C^16,20^. Protein content was measured using bicinchoninic acid protein assay (Pierce BCA Protein Assay Kit, Thermo Fisher Scientific, Waltham, MA, USA 23227). Samples were processed at the Stanford Human Immune Monitoring Center using a mouse 39-plex Affymetrix (San Diego, CA, USA) kit. **Supplementary Table S1** lists all cytokines analyzed.

### Reverse Phase Protein Assay (RPPA)

MFPs from BALB/c mice were resected and processed as described previously^21^. Briefly, tissues were homogenized, lysed, and stored at -80°C until being processed at the Functional Proteomics RPPA Core Facility at The University of Texas MD Anderson Cancer Center. **Supplementary Table S2** lists all assayed proteins.

### Immunohistochemistry (IHC)

Tissues were removed from mice and placed in 10% formalin for 24 hours at 4°Cand then in 70% ethanol before embedding in paraffin and sectioning. Sections (4 µm) were deparaffinized in xylene, rehydrated, boiled in citric acid (10 mM, pH 6) for antigen retrieval, and treated with 3% hydrogen peroxide. Blocking in 10% goat serum was followed by incubation overnight at 4°C CD4 (1:100; eBioscience, San Diego, CA, USA MA1-146) primary antibodies. Sections were incubated with biotinylated secondary antibodies (Vector Laboratories, Newark, CA, USA SK4285) followed by incubation with the substrate using the DAB Peroxidase substrate Kit (Vector Laboratories SK4105) and then counterstained with hematoxylin. A corresponding no primary antibody control was performed for all conditions to confirm specificity. Samples were imaged using an inverted Leica (Wetzlar, Germany) DMi8 microscope.

### SDS-PAGE and Western Blotting

Cell lysates were collected from cells plated in 10 cm dishes, washed twice with ice-cold PBS, and incubated with RIPA Buffer (Sigma-Aldrich, St. Louis, MO, USA R0278) containing 5 mM EDTA (Corning, Corning, NY, USA 46-034-CI), 1 EDTA-free mini cOmplete EASYpack tablet tabled (Roche, Basel, Switzerland 04693159001), and 1 mini PhoSTOP EASYpack (Roche 04906845001). Following mechanical disruption in lysis buffer, supernatant was collected, incubated on ice, sonicated twice (Fisher Scientific, Hampton, NH, USA Sonic Dismembrator Model 100), and spun down at 4°C at 13,300xg. Supernatant was collected and stored at -80°C until use. 10 µg of protein extract, determined by the BCA Protein Assay Kit (Pierce BCA Protein Assay Kit, Thermo Fisher Scientific 23227), was diluted in RIPA buffer and combined 3:1 with 4X sample loading buffer (LI-COR, Lincoln, NE, USA 928–40004) containing 355 mM 2-mercaptoethanol. Samples were vortexed and heated at 95°C for 5 min.

Samples were then loaded onto a 10% TRIS-glycine sodium dodecyl sulfate (SDS) polyacrylamide gels along with Precision Plus Protein Kaleidoscope Prestained Protein Standard (Bio-Rad, Hercules, CA, USA 1610375). Resolving gels were formulated with 30% acrylamide mix (BioRad), 1.5M Tris-HCI pH 8.8 resolving gel buffer, 10% SDS, 10% ammonium persulfate, and TEMED (BioRad) and prepared according to previously established protocols^22^. 5% stacking gels were formulated with DI water, 30% acrylamide mix, 1.5M Tris-HCI pH 6.8 stacking gel buffer (BioRad), 10% SDS, 10% ammonium persulfate, and TEMED^22^. Gel electrophoresis was performed in buffer containing 3.03 g/L TRIS base (Research Products International [RPI], Mount Prospect, IL, USA T60040), 1.0g/L SDS (RPI L22010), and 14.4 g/L glycine (RPI G36050) by supplying 250 V for 25–30 min at room temperature using a Bio-Rad PowerPac HC and mini-PROTEAN Tetra system.

Polyvinylidene difluoride (PVDF) membranes were activated with methanol before washing in ice-cold transfer buffer for 5 minutes. Protein was transferred to a 0.45 μm pore size PVDF membrane using transfer buffer containing 3.03 g/L TRIS base, 14.4 g/L glycine, and 20% methanol by supplying 34 V for 16 h at 4°C. Following transfer, membranes were dried for 10 min at 37°C then incubated in methanol for 30 s for re-activation and washed with TBS for 5 min. Membranes were subsequently blocked for 1 h at room temperature using Intercept TBS Blocking Buffer (LI-COR 927–60001). Blocked membranes were incubated in primary antibodies for target protein and loading control diluted with Intercept TBS buffer and 0.2% TWEEN 20 (Sigma-Aldrich P1379) at the following concentrations: Phospho-p38 MAPK alpha (Thr 180, Tyr182; 1:1000, Invitrogen MA515177), phospho-NF-κB p65 (Ser536; 1:1000, Cell Signaling, Danvers, MA, USA 3033S), and rat alpha-tubulin (1:1000, Invitrogen MA1-80017). Membranes were washed 4x, 5 min each, using TBS containing 0.2% Tween 20 (Invitrogen) and then incubated donkey anti-rabbit IRDye 800CW (1:15000, LICOR 926–32213) or goat anti-rat IRDye 680RD (1:15000, LICOR 926–68076). Membranes were imaged on the Odyssey Fc imager (LICOR) in 600, 700, and 800 nm channels with 30 sec or 2 min exposure time, as necessary. Data were analyzed to determine protein expression relative to loading controls using Image Studio v6.0 (LI-COR Biosciences).

### 3D Growth Assay

To determine the effects of Th2-derived factors on cancer cells, a 3D invasion assay was performed where 4T1s were plated in 100uL of 2.5mg/ml Matrigel (Corning 354230) at 2500 cells/gel in a flat bottom 96 well plate. Media with and without TGFβ1 (10 ng/mL, Peprotech 100-21) was added on top of the gel. The media was changed daily. Brightfield images were taken of cells colonies within gels. Images were then analyzed in FIJI where buds emerging from colonies were counted^23^.

### Quantitative Real Time PCR (qPCR)

Cells were harvested by centrifugation and washed once with cold PBS. Cell pellets were snap-frozen and stored at −80°C until RNA extraction. For isolation, frozen pellets were thawed on ice and lysed directly in PureLink™ RNA Mini Kit lysis buffer (Thermo Fisher Scientific 12183025) supplemented with β-mercaptoethanol. Lysates were homogenized by pipetting and applied to spin columns according to the manufacturer’s instructions. Columns were washed and total RNA was eluted in RNase-free water. RNA concentration and purity were determined using a NanoDrop spectrophotometer (Thermofisher). Samples with A260/280 ratios between 1.9 and 2.1 were used for downstream applications. Complementary DNA (cDNA) was synthesized from 500 ng–1 µg of total RNA using the High-Capacity cDNA Reverse Transcription Kit (Applied Biosystems, Foster City, CA, USA Thermo Fisher Scientific 4368814) according to the manufacturer’s protocol. Reverse transcription reactions (20 µL total volume) were performed under the following conditions: 25°C for 10 minutes, 37°C for 120 minutes, 85°C for 5 minutes, and hold at 4°C. cDNA was diluted 1:5 with nuclease-free water prior to quantitative PCR. qPCR was performed using PowerUp™ SYBR™ Green Master Mix (Thermo Fisher Scientific, A25742) on a QuantStudio™ 5 Real-Time PCR System (Applied Biosystems). Reactions were performed in triplicate in 20 µL volumes containing 2× SYBR Green Master Mix, 200 nM forward and reverse primers, and 2 µL diluted cDNA. Thermal cycling conditions were: 95°C for 2 minutes, 40 cycles of 95°C for 15 seconds, and 60°C for 30 seconds. Melt curves were analyzed to confirm specificity. No-template and no-reverse transcriptase controls were included. Relative gene expression was calculated using the ΔΔCt method and normalized to β-actin. The primers used for qPCR are listed in **Supplementary Table S3**.

### Transwell Assays

Invasion/migration assays were performed with 4T1 and BMDMs using CM from Th2 cells as a chemoattractant (Corning 354483 & 354578). 4T1s or BMDMs were seeded at 2×10^5^ cells/mL on the top on the insert. Basal media was used as a negative control, and test conditions were media spiked TGFβ (10 ng/mL, Peprotech 100-21), Th2 CM, and Th2 CM with added aTGFβ (2.5 µg/mL, R&D, Minneapolis, MN, USA MAB1835) neutralizing antibody. Cells that invaded through the Matrigel inserts were stained with ProLong Glass Antifade Mountant with NucBlue Stain (Invitrogen P36981) and counted.

### Statistical Analysis

Data are presented as mean ± SEM. Statistical analyses were performed using GraphPad Prism 10 (GraphPad Software, San Diego, CA, USA). Comparisons between two groups were conducted using unpaired two-tailed t-tests. Comparisons among three or more groups were analyzed using one-way ANOVA followed by Tukey’s multiple comparison post-hoc test. For experiments involving two independent variables, two-way ANOVA was used where appropriate.

## Results

### Immunodeficient Irradiated Tissue Accumulates CD4+ T-cells

To determine how immune composition shapes the post-irradiation (post-IR) microenvironment, we quantified CD4+ T-cell infiltration in irradiated mammary fat pads (IR-MFP) under immunocompetent and CD8+ T-cell-depleted (immunodeficient) conditions (**Figure 1A**). Immunodeficient mice experience significantly greater CD4+ T-cell infiltration to the IR-MFP (**Figure 1B**) as compared to immunocompetent controls. After 10 days, the timepoint at which tumor cell infiltration is noted^9^, infiltration decreases but immunodeficient mice maintain higher CD4+ T-cell infiltration (**Figure 1C**), indicating a sustained shift in immune composition in the absence of CD8+ T-cell–mediated regulation. Importantly, the timing of CD4+ T-cell infiltration coincides with macrophage recruitment as previously reported^9^, peaking at day 5. This coordinated timing suggests that CD4+ T-cells may contribute to the establishment of a macrophage-supportive microenvironment following radiation. Given poor outcomes for patients with a high CD4/CD8 ratio^14^ and the crosstalk between CD4+ T-cells and macrophages^13^, this raises the possibility that CD4+ T-cell enrichment promotes macrophage accumulation in this setting. Notably, macrophage and CD4+ T-cell infiltration followed by eventual tumor infiltration at day 10 for IR-immunodeficient mice suggests a sequential model in which CD4+ T-cells contribute to a pro-tumor niche. These findings establish that irradiated immunodeficient tissue accumulates CD4+ T-cells coincident with macrophage recruitment, creating conditions permissive for tumor cell recruitment.

**Figure 1.**
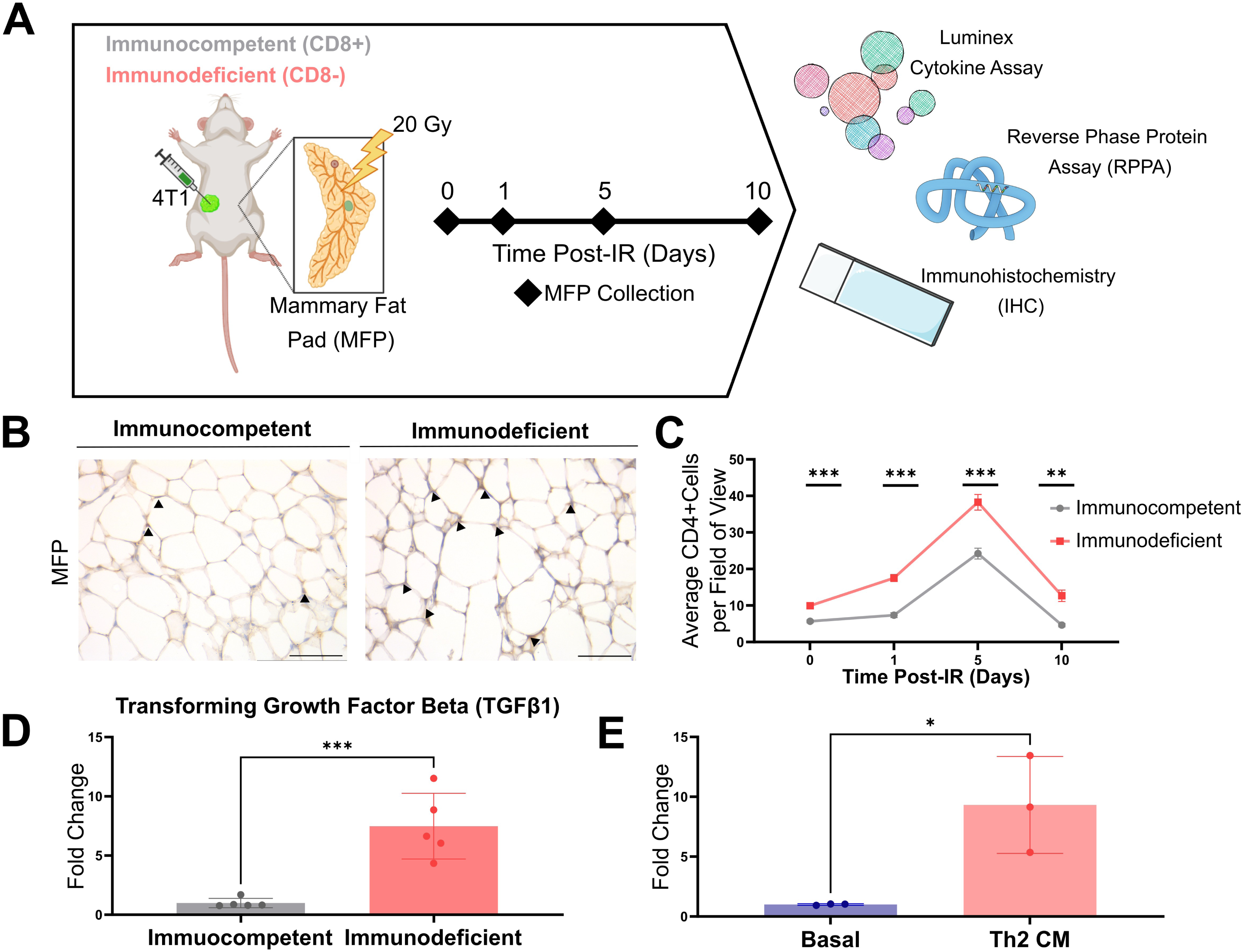
CD4+ T-cells infiltrate irradiated tissues in immunodeficient mice. (A) *In vivo* experimental schematic. Immunocompetent (IgG) or immunodeficient (aCD8) mice (n=5 mice/condition) inoculated with a 4T1 TNBC tumor were irradiated to a dose of 20 Gy after tumors reached 100mm^3^, and mammary fat pads (MFPs) were harvested after 0, 1, 5, and 10 days. (B) Immunohistochemistry (IHC) was performed to detect CD4+ T-cell infiltration into MFPs, and representative images at day 5 area shown. Black arrows indicate CD4+ T-cells. The corresponding quantification in (C) revelaed increased CD4+ staining in immunodeficient mice. Scale bar is 100 µm. (D) Transforming growth factor beta 1 (TGFβ1) was increased in the MFP of irradiated immunodeficient mice compared to unirradiated controls as determined by a Luminex immunoassay. (E) Cytokine array (n=3 independent replicates) indicated the presence of TGFβ1 in Th2 conditioned media (CM) isolated from mice and polarized *in vitro* at nearly 10-fold the amount as compared to basal media. Error bars show standard error. Statistical significance was determined by an unpaired two tailed t-test with *p<0.05, **p<0.01, and ***p<0.001.

### Th2 Conditioned Media (CM) Promotes TNBC and Macrophage Invasion

Given the accumulation of CD4+ T-cells in irradiated immunodeficient tissue, we next asked whether CD4+ T-cell–derived signals functionally contribute to the pro-tumor microenvironment. We focused on Th2-polarized CD4+ T-cells, which are also responsible for the cytokine profile that polarizes macrophages to the immunosuppressive M2 phenoptye^24^, are associated with immunosuppression, recurrence, and poorer outcomes for patients^13,25^. Understanding the difference in the secretome between immunocompetent and immunodeficient mice is important in determining how immunodeficient mice develop a niche where TNBC cells can colonize tissues. To characterize the cytokine milieu present at the IR-MFP, we used a Luminex cytokine assay performed on the whole MFP homogenate and observed over 7-fold upregulation of transforming growth factor beta (TGFβ) (**Figure 1D**). To study the influence of TGFβ on tumor cell invasion in 3D, 4T1 cells embedded in Matrigel were treated with TGFβ, and an increase in 4T1 colony budding was observed (**Supplementary Figure S1A,B**). Th2 CM also contained detectable levels of TGFβ1 (**Figure 1E**), suggesting that these cells may contribute directly to the pro-tumor cytokine milieu post-IR.

Functionally, Th2 CM also enhanced 4T1 TNBC and bone marrow-derived macrophage (BMDM) invasion (**Figure 2A,B**). Neutralization of TGFβ1 significantly abrogated this effect in both cell types. While TGFβ1 is already an established regulator of cancer invasiveness^26^, these findings implicate Th2 cells as a functional source of TGFβ1, which promotes macrophage and tumor cell recruitment. This is further supported by western blot results showing that Th2 CM increases invasive marker vimentin expression of 4T1 cells in a TGFβ1-dependent manner (**Supplementary Figure S1C**). Together, these data suggest a functional link between CD4+ T-cell accumulation and a pro-invasive microenvironment with Th2-derived TGFβ1 as a key mediator of this transition.

**Figure 2.**
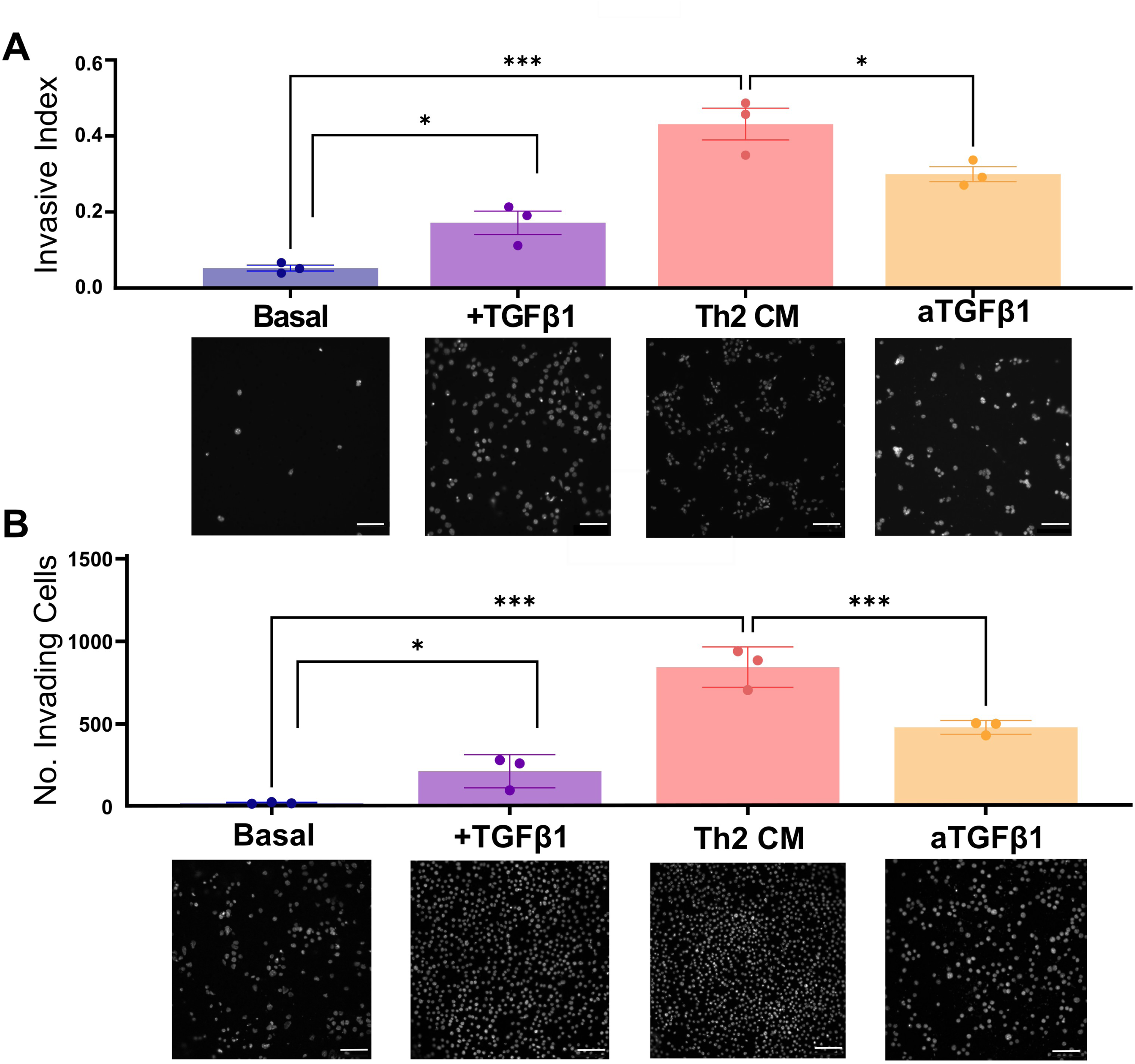
The Th2 secretome increases invasive behavior of TNBC cells and macrophages. 4T1 TNBC cells (A) and bone marrow derived macrophages (BMDMs) (B) incubated with basal media containing recombinant TGFβ1 or Th2 CM increased invasion compared to basal media (n=3 independent replicates). Neutralizing TGFβ1 in Th2 CM (aTGFβ1) partially abrogated this invasive response. Error bars show standard error. Statistical significance was determined by one-way ANOVA analysis with *p<0.05 and ***p<0.001.

### Th2-Derived TGF**β**1 Induces a Feed-Forward Amplification Loop

Having identified the TGFβ1-Th2 relationship capable of increasing tumor cell invasion, we sought to elucidate the regulatory signaling pathways. qPCR showed significant upregulation of *Tgfb1* in both 4T1 TNBC (**Figure 3A**) cells and primary BMDMs (**Figure 3B**) following Th2 CM exposure, indicating that the Th2 secretome is sufficient to induce endogenous TGFβ1 production by 4T1 TNBC cells and BMDMs. Neutralizing Th2 CM TGFβ1 significantly reduces *Tgfb1* expression, suggesting that TGFβ signaling is required to sustain this amplification loop. BMDMs were also polarized toward an M2 phenotype through increased *Mrc1* and reduced *Nos2* expression following incubation with Th2 CM (**Figure 3C**). This demonstrates that exogenous TGFβ1 from Th2 cells drives the production of endogenous TGFβ1 at the treated site, which enhances immunosuppressive signaling in IR-MFPs. These findings support a model in which Th2-derived TGFβ1 initiate a self-renewing feed-forward loop which sustains and increases TGFβ1 levels in the IR-MFP in the absence of CD8+ T-cells. This mechanism would progressively enrich the TGFβ1 levels at the site, as we observed with TGFβ1 levels increasing up over the course of 10 days, reinforcing the immunosuppressive, macrophage-laden, tumor-permissive niche.

**Figure 3.**
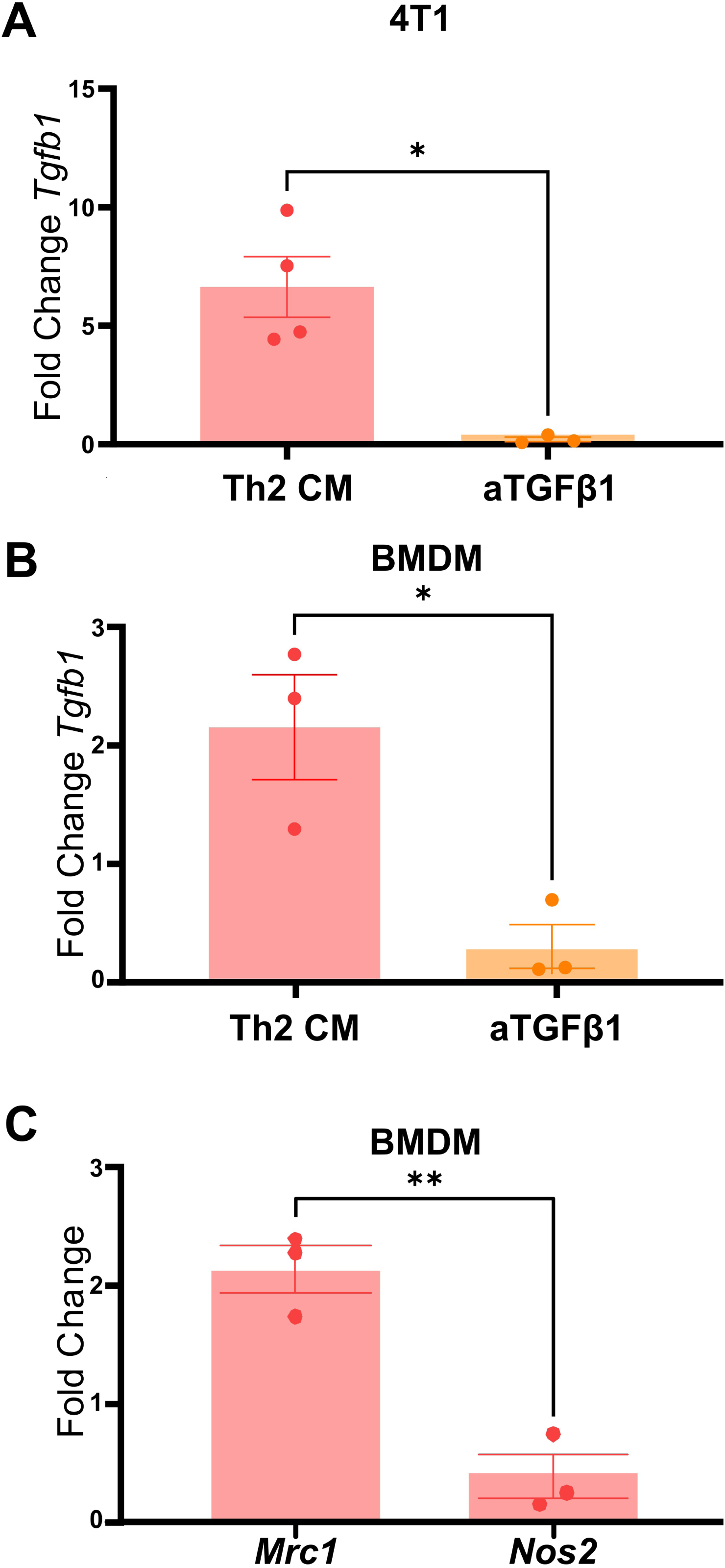
The Th2-driven increase in *Tgfb1* expression in TNBC cells and macrophages is dependent on secreted TGFβ1. 4T1 TNBC cells (A) and BMDMs (B) increase expression of *Tgfb1* when treated with Th2 CM with respect to basal media. This effect is abrogated when TGFβ1 is neutralized (n=3 independent replicates). BMDMs (C) increase *Mrc1*, the gene encoding CD206, and decrease Nos2, the gene encoding iNOS, when treated with Th2 CM with respect to basal media. Rat α-tubulin was used as a housekeeping gene (n= 3 independent replicates). Error bars show standard error. Statistical signifiance determined by one-way ANOVA analysis with *p<0.05 and **p<0.01.

### Th2 CM Activates Non-Canonical TGF**β** Signaling in TNBC Cells

To understand the downstream signaling pathways that the Th2-TGFβ1 axis engages, we performed reverse phase protein assay (RPPA) on resected MFPs. Despite increased TGFβ1 levels, the immunodeficient IR-MFP shows no significant changes in pSMAD3 expression, a key player in canonical TGFβ1 signaling (**Supplemental Figure S2A**). In contrast, increased p38 MAPK, a downstream proteins in non-canonical TGFβ1 signaling, is observed (**Figure 4A**). This indicates that TGFβ1 signaling in this context preferentially engages non-canonical pathways.

**Figure 4.**
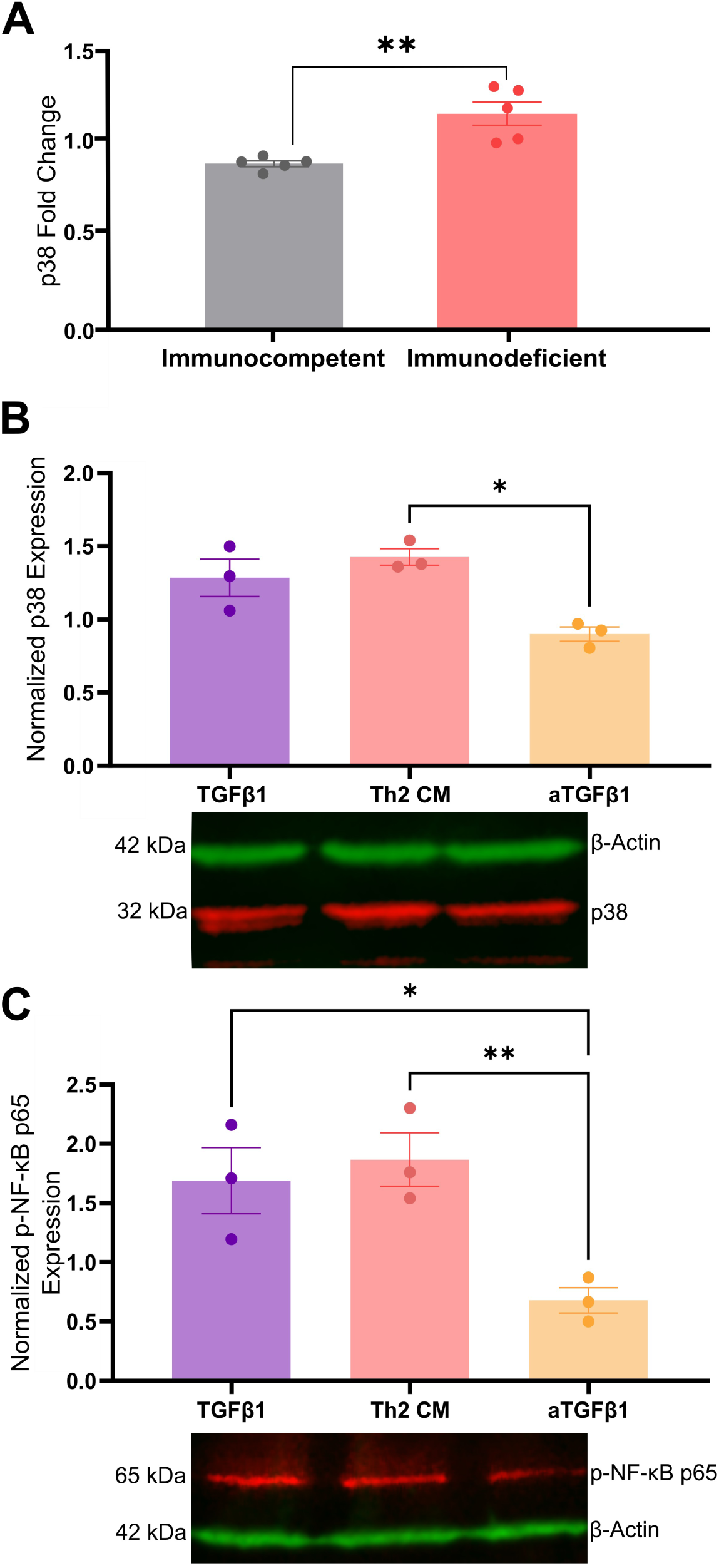
The Th2 secretome signals through p38 and NF-κB. (A) Protein expression of p38 is elevated in the MFP of irradiated immunodeficient mice as determined by RPPA analysis (n=5 mice/condition). Statistical significance was determined by an unpaired two tailed t-test with **p<0.01. Quantification of relative protein expression of p38 (B) and NF-κB phosphorylation (C) in 4T1 tumor cells with respect to basal media with β-actin as loading control and representative western blot shown below (n=3 independent replicates). Adding recombinant TGFβ1 to basal medium or Th2 CM increased expression while TGFβ1 neutralization (aTGFβ1) in the Th2 CM reduced expression of both. Error bars show standard error. Statistical significance determined by one-way ANOVA analysis with *p<0.05 and **p<0.01.

In agreement with *in vivo* findings, western blotting of 4T1 TNBC cells treated with Th2 CM show increased p38 MAPK expression (**Figure 4B**) while p-SMAD2/3 is unchanged by the Th2 secretome (**Supplemental Figure S2B**). These findings suggest that the Th2 secretome may bias TGFβ1 toward non-canonical signaling in tumor cells. Further analysis revealed that Th2 CM increases TNBC cell NF-κB phosphorylation, which is attenuated by antibodies against TGFβ1 (**Figure 4C**). In contrast, BMDMs did not exhibit significant activation of p38 or NF-κB in response to Th2 CM (**Supplemental Figure S3**). Taken together, these findings indicate that Th2-derived TGFβ effect is cell-specific and engages non-canonical p38/NF-κB signaling in TNBC cells to promote invasive behavior. This provides a mechanistic basis for the pro-tumor niche that emerges in the immunodeficient irradiated microenvironment.

## Discussion

Immunodeficiency creates an immune landscape in which Th2 CD4+ T-cells accumulate and reshape irradiated normal tissue into a TGFβ-rich, pro-invasive niche (**Figure 5**). Mechanistically, our data support a TGFβ1-dependent feedback loop where Th2 cell-derived TGFβ1 drives the invasive behavior of both TNBC cells and macrophages. TNBC cells and macrophages respond to Th2 CM by producing additional TGFβ1, establishing a feed-forward loop that reinforces invasion and immunosuppression. Macrophages maintained an M2-like phenotype, which is consistent with their anti-inflammatory, tumor-supportive role. Simultaneously, Th2 cells increased TNBC cell survival and invasion by activating p38 MAPK and NF-κB in TNBC cells while pSMAD2/3 remained unchanged, indicating non-canonical TGFβ signaling^27^. Increased survival of recruited TNBC cells likely contributes to their re-establishment and the progression of recurrence.

**Figure 5.**
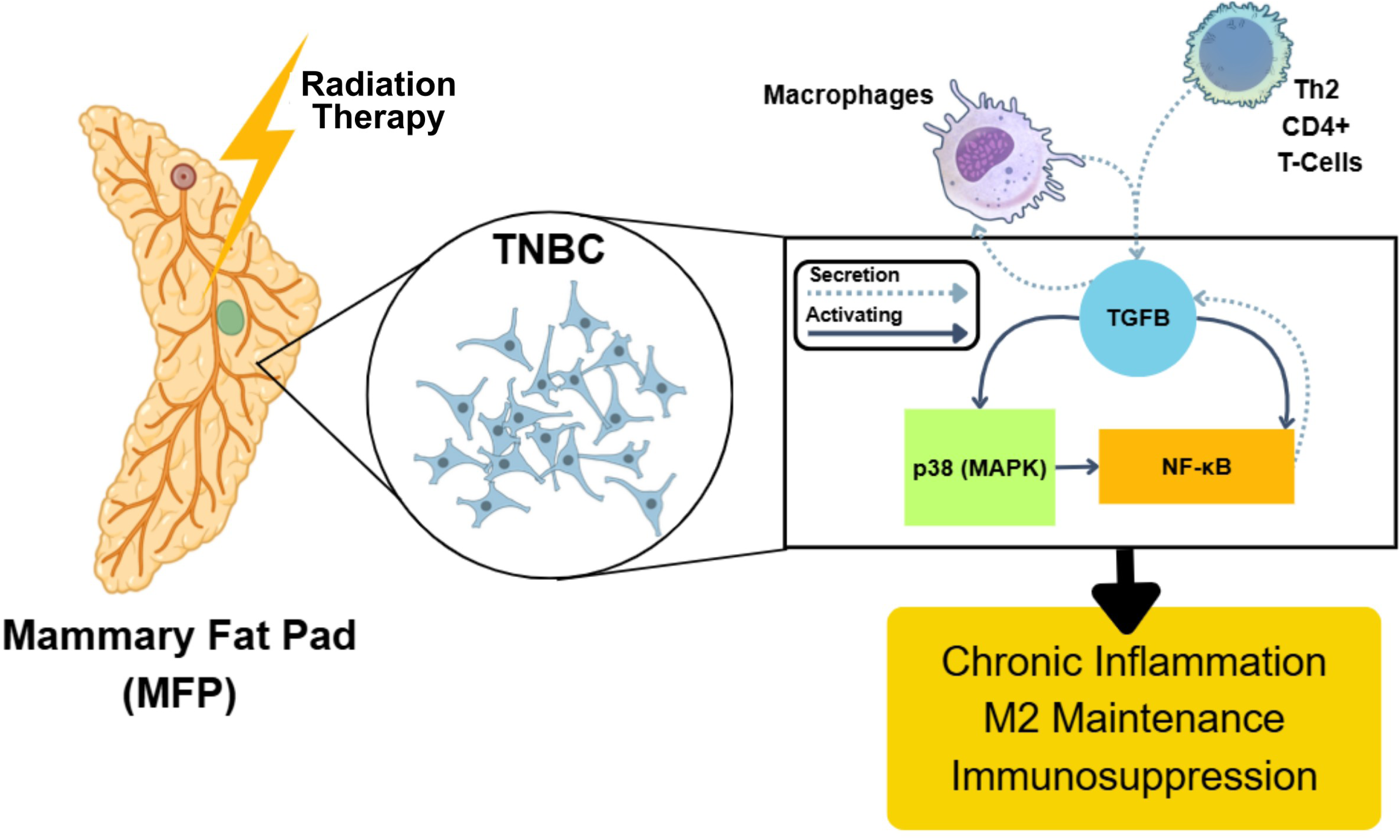
Model of Th2-derived TGFβ1 driving a positive, feed-forward loop in recurrence post-therapy. This loop leads to increased p38 MAPK and NF-κB activation in 4T1 TNBC cells, enforcing macrophage M2 polarization, tumor cell invasiveness, and inflammation.

As p38 MAPK is a stress responder, its activation post-IR with TGFβ1 is expected^28^. However, its role in partnership with Th2-derived TGFβ is unique. This agrees with findings that p38 is vital for breast cancer metastasis and resistance to immunotherapy^29^. NF-κB is similarly activated after radiation damage where it leads to macrophage recruitment and immunosuppression^30,31^, which we also observed in our model. Completing the loop, the p38 MAPK/NF-κB pathway is well-documented as a driver of cancer metastasis^32^ and growth^33,34^. In contrast, macrophages exhibit minimal activation of these pathways, suggesting that TGFβ1 promotes macrophage invasion through mechanisms distinct from those observed in TNBC cells. Th2 CM-induced activation of NF-κB in TNBC cells positions it as a central hub integrating TGFβ1 signaling, linking cytokine exposure to functional invasion outcomes.

TGFβ1 is well-recognized for its roles in tumor progression^35,36^, immunosuppression^37,38^, and treatment resistance^39,40^. TGFβ has been shown to facilitate the invasive behavior of cancer cells, but it remains poorly understood in the context of the normal tissue. Though dysregulated SMAD signaling is closely associated with tumor growth and epithelial-mesenchymal transition^41^, there are other TGFβ1 targets capable of pro-tumor signaling. The noncanonical TGFβ1 pathway exhibits diverse signaling and downstream activation such as NF-κB and p38 MAPK^27^. Here, we find a role for tumor-permissive, non-canonical TGFβ signaling post-radiation which increases the vulnerability of immunodeficient patients to recurrence.

The immune microenvironment following radiation is highly dynamic, shaped by lymphocyte depletion, stromal injury, and compensatory immune repopulation^42–48^. While CD4+ T-cell infiltration can associate with either tumor control^49–51^ or progression^11,37,52^, our findings suggest that Th2 polarization skews this balance toward tissue remodeling and tumor permissiveness within irradiated tissue, in agreement with similar findings^13,25,53^. Rather than function through classical effects, Th2 cells appear in this site as early orchestrators, enriching the tissue with TGFβ and promoting macrophage-mediated amplification. This context-dependent behavior crystallizes the importance of immune composition, not just presence, in determining post-therapy outcomes.

Cytokine production in irradiated tissue arises from multiple cellular sources, and while Th2 cells contribute to TGFβ1 enrichment, additional stromal and immune populations likely participate. Performing cytokine Luminex and RPPA analyses on whole MFP lysates limited our ability to directly pinpoint the cell types that regulate TGFβ levels. In addition, while we observe that Th2 cells facilitate a tissue microenvironment conducive to recurrence, it is possible that another CD4+ T-cell subtype is involved. Future studies defining CD4+ T-cell subset dynamics in irradiated tissue will further clarify these interactions. Finally, although TGFβ neutralization partially reduced invasion, the residual activity likely reflects cooperative signaling within the Th2 secretome in agreement with previous findings^54–56^, underscoring the complexity of immune-driven microenvironmental remodeling.

Overall, these findings identify a TGFβ-dependent amplification circuit in which Th2 cells enrich irradiated tissue with TGFβ and induce macrophage *Tgfb1* expression under immunocompromised conditions, sustaining a cytokine-rich niche. While macrophages amplify this environment, non-canonical p38 MAPK and NF-κB activation in TNBC cells directly links Th2-derived TGFβ to tumor cell invasion. These data shift attention from tumor-intrinsic resistance to irradiated normal tissue as an active driver of recurrence and suggest that selective interruption of the Th2–TGFβ axis may mitigate TNBC relapse following radiotherapy, improving outcomes for breast cancer patients. Complete understanding of the pro-tumor immune milieu will allow for early identification of high-risk patients and personalized approaches to mediating the tissue immune ecosystem to prevent recurrence.

## Additional Information

### CRediT Authorship Contribution Statement

**McKenzie A. Mayeaux:** Conceptualization, Methodology, Investigation, Validation, Formal Analysis, Writing – Original Draft, Writing – Review & Editing, Visualization. **Benjamin P. Altman:** Investigation. **Benjamin C. Hacker:** Investigation. **Steven M. Alves:** Investigation. **Dadi Jiang:** Methodology, Formal Analysis. **Albert C. Koong:** Project Administration. **Edward E. Graves:** Conceptualization. **Marjan Rafat:** Conceptualization, Methodology, Investigation, Resources, Funding Acquisition, Writing – Review & Editing, Supervision, Project Administration.

## Acknowledgments

We thank Dr. Christopher Contag for luciferase-labeled 4T1s, Dr. Rachelle Johnson for use of QuantStudio 5, Dr. Jamey D. Young for use of the homogenizer, and Dr. Heather Pua for assistance in establishing primary T-cell cultures. We thank the Translational Pathology Shared Resource core facility for *ex vivo* sample preparation (P30CA068485). Data were generated in part through the use of the Functional Proteomics Reverse Phase Protein Array Core (RPPA), which receives partial support from the National Cancer Institute under grant P30CA016672 to UT MD Anderson and Dr. Yiling Lu’s NIH R50 Grant #R50CA221675. The research reported in this publication was not directly funded through the grant P30CA016672 to The University of Texas MD Anderson Cancer Center and is not within the scope of such grant. The schematics were prepared using BioRender, the NIH BioArt resource, and the MIT BioIcons resource.

## Data Availability

The datasets used and/or analyzed for the present work are available from the corresponding author on reasonable request.

## Ethics Statement

Animal experiments were performed in accordance with ARRIVE 2.0 guidelines and approved by the Vanderbilt University Institutional Animal Care and Use Committee (approval no. M1800017-02).

