## Supplementary Figures for "CD4+ T-Cells Drive Triple Negative Breast Cancer Recurrence via Non-Canonical TGFβ Signaling"

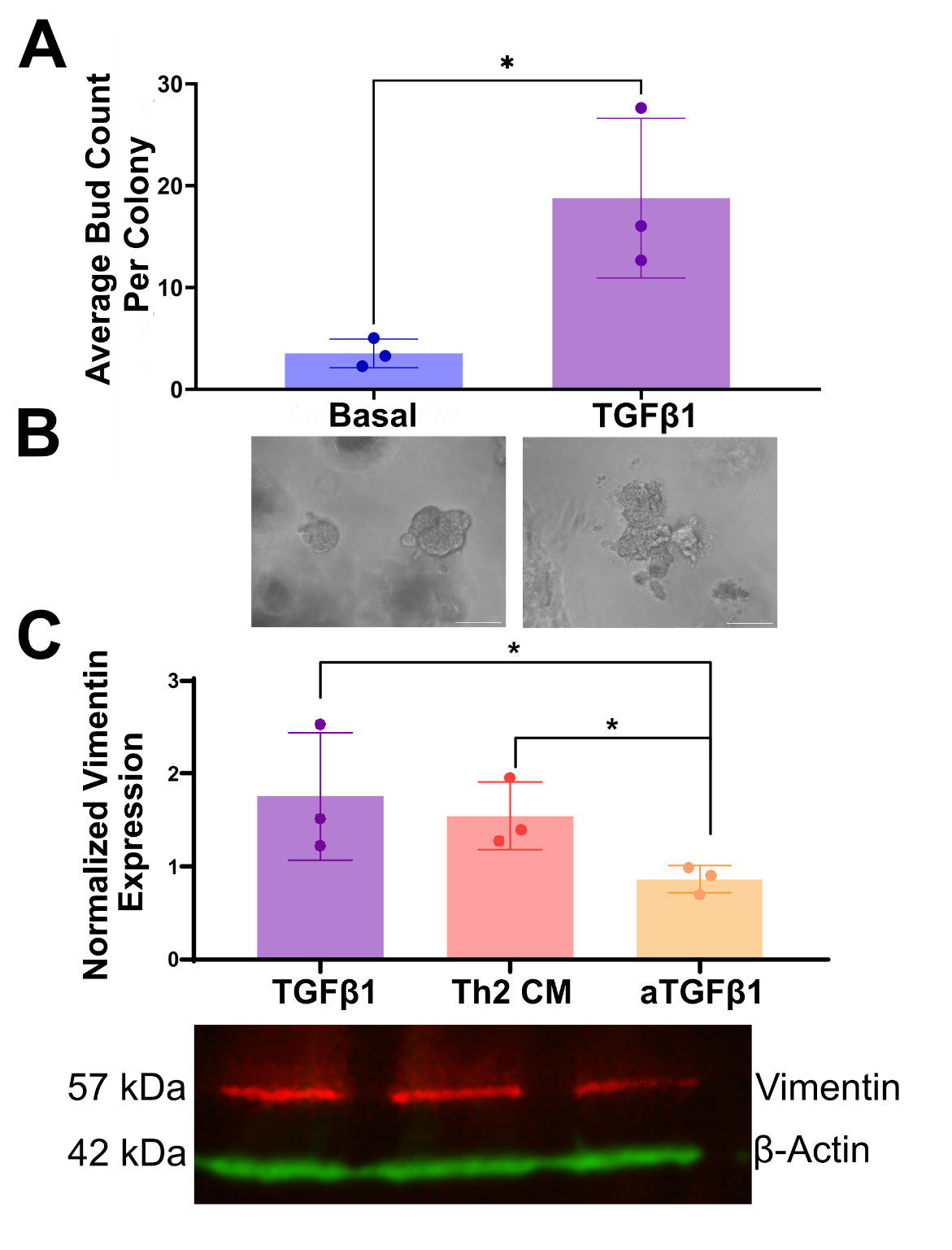

Figure S1. TGFβ1 increases budding and vimentin expression in 4T1 TNBC cells. (A) Colony budding of 4T1 TNBC cells embedded in 2.5 mg/ml Matrigel increased significantly with TGFβ1 (n=3 independent replicates). Representative images shown in (B). Scale bars are 100 μm. Statistical significance determined by unpaired two-tailed student’s t-test with *p<0.05. (C) 4T1 TNBC cells increased vimentin expression when incubated with basal media containing recombinant TGFβ1 or with Th2 conditioned media (CM). Data normalized to expression in basal media. Neutralizing TGFβ1 in Th2 CM (aTGFβ1) reduced this invasive response (n=3 independent replicates). Error bars show standard error. Statistical significance was determined by one-way ANOVA analysis with *p<0.05.

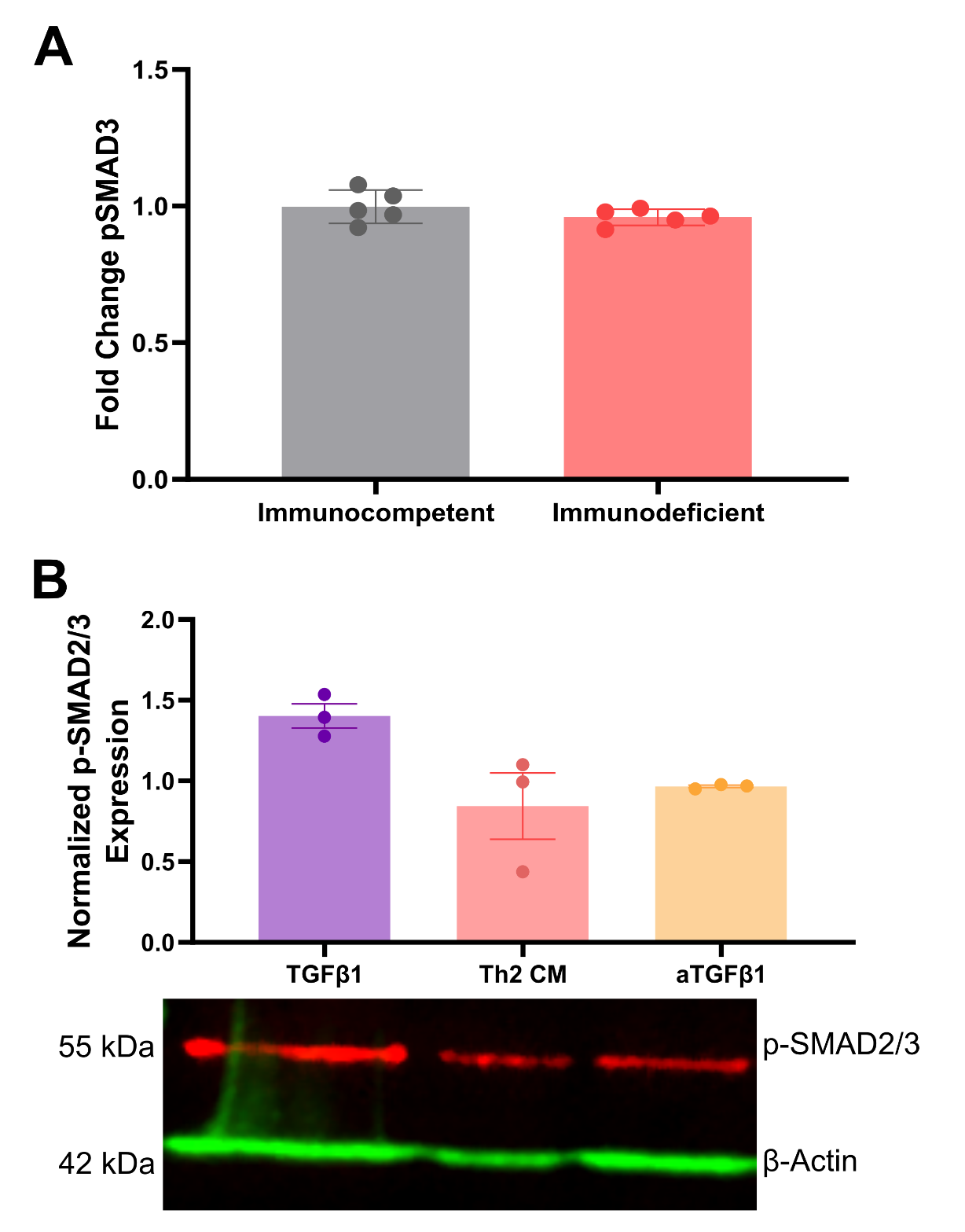

**Figure S2. TGFβ1 does not signal through the canonical SMAD pathway *in vivo* or in response to Th2 CM.** (A) Protein expression of pSMAD3 is unaffected in the mammary fat pad of irradiated, immunodeficient mice as determined by RPPA analysis. Data normalized to unirradiated, immunocompetent mice (n=5 mice/condition). Statistical significance was determined by an unpaired two tailed t-test. (B) Quantification of relative protein expression of pSMAD2/3 in 4T1s with β-actin as loading control (n=3 independent replicates). Error bars show standard error. Statistical significance determined by one-way ANOVA analysis.

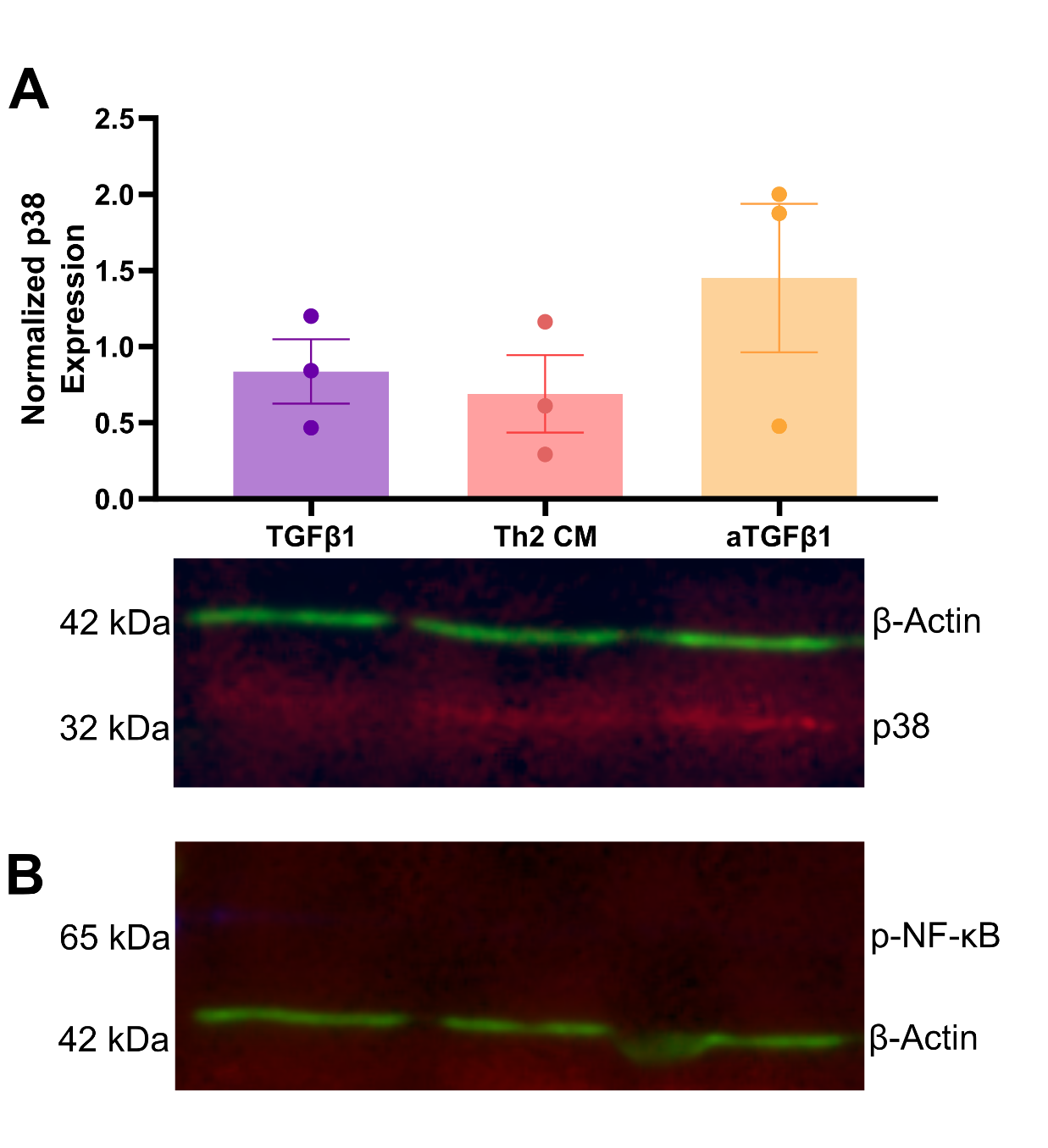

**Figure S3. Macrophages do not signal through p38 or p-NF-**κ**B p65 in response to Th2 CM.** Quantification of relative protein expression of (A) p38 and (B) p-NF-κB p65 in bone marrow-derived macrophages incubated with basal media containing recombinant TGFβ1 or with Th2 CM. β-actin used as loading control (n=3 independent replicates). Expression normalized to cells in basal media. Error bars show standard error. Statistical significance determined by one-way ANOVA analysis.

**Table S1. List of cytokines evaluated in immunodeficient mice 10 days post-IR using a 39-plex immunoassay.**

| Fold Change >1.5 | |
| --- | --- |
| Cytokine Name | **Abbreviation** |
| Interleukin-23 | IL-23 |
| Interleukin-6 | IL-6 |
| Transforming growth factor beta | TGFβ |
| Tumor necrosis factor alpha | TNFA |
| Fold Change <1.5 | |
| Cytokine Name | **Abbreviation** |
| Interleukin-2 | IL-2 |
| Eotaxin (C-C Motif chemokine ligand 11) | CCL11 |
| Granulocyte-Macrophage Colony-Stimulating Factor | GM-CSF |
| Gro-alpha/Chemokine (C-X-C motif) ligand 1 | GROA/CXCL1 |
| Granulocyte colony stimulating factor/colony stimulating factor 3 | G-CSF/CSF3 |
| Interferon alpha | IFNα |
| Interferon gamma | IFNγ |
| Interleukin-10 | IL-10 |
| Interleukin-12 p70 | IL-12p70 |
| Interleukin-13 | IL-13 |
| Interleukin-15/Interleukin-15 Receptor | IL-15/IL-15R |
| Interleukin-17A | IL-17A |
| Interleukin-18 | IL-18 |
| Interleukin-1 alpha | IL-1α |
| Interleukin-1 beta | IL-1β |
| Interleukin-22 | IL-22 |
| Interleukin-27 | IL-27 |
| Interleukin-28 | IL-28 |
| Interleukin-3 | IL-3 |
| Interleukin-31 | IL-31 |
| Interleukin-4 | IL-4 |
| Interleukin-5 | IL-5 |
| Interleukin-9 | IL-9 |
| Interferon γ-induced protein 10 kDa/ Chemokine (C-X-C motif) ligand 10 | IP-10/CXCL10 |
| Leptin | LEPTIN |
| Leukemia inhibitory factor | LIF |
| Lipopolysaccharide-induced CXC chemokine/ Chemokine (C-X-C Motif) ligand 5 | LIX |
| Monocyte chemoattractant protein-1/ C-C Motif chemokine ligand 2 | MCP1/CCL2 |
| Monocyte chemoattractant protein-3/ C-C Motif chemokine ligand 7 | MCP3/CCL7 |
| Macrophage colony-stimulating factor | M-CSF |
| Macrophage inflammatory protein-1 alpha/ C-C Motif chemokine ligand 3 | MIP1α/CCL3 |
| Macrophage inflammatory protein-1 beta/ C-C Motif chemokine ligand 4 | MIP1β/CCL4 |
| Macrophage inflammatory protein-2/ Chemokine (C-X-C motif) ligand 2 | MIP2/CXCL2 |
| Regulated on activation, normal T cell expressed and secreted/C-C Motif Chemokine Ligand 5 | RANTES/CCL5 |
| Vascular Endothelial Growth Factor | VEGF |

**Table S2. List of proteins evaluated in immunodeficient mice 10 days post-IR using reverse phase protein assay (RPPA).**

| Fold Change >1.2 | | | | |
| --- | --- | --- | --- | --- |
| GYS1 | INSR | H3 | NF2 | HSPB1 pS82 |
| AXL | RAD51 | STAT3 pY705 | NDRG1 pT346 | STMN1 |
| Fold Change <1.2 | | | | |
| 14-3-3ζ | 14-3-3β | 4E-BP1 | 4E-BP1 pS65 | 53BP1 |
| ARAF | ACC1 | ACC1 pS79 | AKT | AKT pS473 |
| AKT pT308 | AMPKα2 pS345 | AMPKα | AMPKα pT172 | AR |
| ARID1A | ATG3 | ATG7 | ATM | ATM pS1981 |
| ATR pS428 | AURKB | BRAF | B7-H4 | BAK |
| BIM | FAK | FAK pY397 | FASN | Fibronectin |
| FOXM1 | FOXO3 | FOXO3 pS318/321 | G6PD | Granzyme B |
| GAB2 | GATA6 | GCLM | GCN5L2 | GLUD1/2 |
| GLS | GZMB | GSK3α/β pS21/S9 | GYS1 pS641 | HER2 pY1248 |
| HER3 | HER3 pY1289 | Heregulin | HES1 | HK2 |
| HSP70 | IGF1R pY1135/1136 | IGFBP2 | IGF1Rβ | INPP4B |
| IRF1 | IRS1 | JAG1 | JAK2 | JNK2 |
| JNK pT183/Y185 | LC3A/B | LCK | LDHA | LRP6 pS1490 |
| MCL1 | MCT4 | MDM2 pS166 | MEK1 | MEK1 pS217/S221 |
| MERIT40 pS29 | MIF | MMP14 | MMP2 | MKNK1 |
| MSH6 | MSI2 | MTOR | MTOR pS2448 | MYH11 |
| MYH9 pS1943 | PKMYT1 | N-Cadherin | Napsin A | NF-κB p65 pS536 |
| NOTCH1 | NOTCH3 | OCT4 | P-Cadherin | p16INK4a |
| p21 | p27Kip1 | p27 pT198 | p38 MAPK | p38 pT180/Y182 |
| ERK1/2 | p53 | p70S6K | p70S6K pT389 | p90RSK pT573 |
| PAICS | PAK1 | PAK4 | PAR | PARP |
| Paxillin | PD-L1 | PDCD4 | PDHK1 | PDK1 |
| PDK1 pS241 | PEA15 | PEA15 pS116 | PI3K p110α | PI3K p85 |
| PKAα | PKCβII pS660 | PKCδ pS664 | PKCα | PKM2 |
| PLCγ2 pY759 | PLK1 | PMS2 | PR | PRAS40 pT246 |
| PREX1 | PTEN | RAB11 | RAB25 | RAD50 |
| Raptor | RBM15 | RB1 pS807/811 | RICTOR | RICTOR pT1135 |
| RIPK1 | RPA32 pS4/S8 | RSK | S6 pS235/236 | S6 pS240/244 |
| SDHA | SHC1 pY317 | SHP2 pY542 | SLC1A5 | SLFN11 |
| SMAD1 | SMAD3 | SOD2 | SOX2 | SRC pY419 |
| SRC pY527 | STAT3 | STAT5A | STING | TAZ |
| TFAM | TFRC | TIGAR | TRIM25 | TSC1 |
| TSC2 | TSC2 pT1462 | TUFM | TYRO3 |  |

**Table S3. qPCR primer sequences.**

|  | Gene | Forward Sequence | Reverse Sequence |
| --- | --- | --- | --- |
|  | *Actb* | CCACCATGTACCCAGGCATT | CGGACTCATCGTACTCCTGC |
|  | *Tgfb* | GTGCGGAAACCCAAACTTTCT | AGAGGTTTGGAGAACCTGCG |
|  | *Mrc1* | GTCAGAACAGACTGCGTGGA | AGGGATCGCCTGTTTTCCAG |
|  | *Nos2* | TGCCAGGGTCACAACTTTACA | CTCTCCACTGCCCCAGTTTT |
